# DPP9 is an endogenous and direct inhibitor of the NLRP1 inflammasome that guards against human auto-inflammatory diseases

**DOI:** 10.1101/260919

**Authors:** Franklin L. Zhong, Kim Robinson, Chrissie Lim, Cassandra R. Harapas, Chien-Hsiung Yu, William Xie, Radoslaw M. Sobota, Veonice Bijin Au, Richard Hopkins, John E. Connolly, Seth Masters, Bruno Reversade

## Abstract

The inflammasome is a critical immune complex that activates IL-1 driven inflammation in response to pathogen- and danger-associated signals. Nod-like receptor protein-1 (NLRP1) is a widely expressed inflammasome sensor. Inherited gain-of-function mutations in NLRP1 cause a spectrum of human Mendelian diseases, including systemic autoimmunity and skin cancer susceptibility. However, its endogenous regulation and its cognate ligands are still unknown. Here we apply a proteomics screen to identify dipeptidyl dipeptidase, DPP9 as a novel interacting partner and a specific endogenous inhibitor of NLRP1 inflammasome in diverse primary cell types from human and mice. DPP9 inhibition via small molecule drugs, targeted mutations in its catalytic site and CRISPR/Cas9-mediated genetic deletion potently and specifically activate the NLRP1 inflammasome leading to pyroptosis and IL-1 processing via ASC and caspase-1. Mechanistically, DPP9 maintains NLRP1 in its monomeric, inactive state by binding to the auto-cleaving FIIND domain. NLRP1-FIIND is a self-sufficient DPP9 binding module and its disruption by a single missense mutation abrogates DPP9 binding and explains the aberrant inflammasome activation in NAIAD patients with arthritis and dyskeratosis. These findings uncover a unique peptidase enzyme-based mechanism of inflammasome regulation, and suggest that the DPP9-NLRP1 complex could be broadly involved in human inflammatory disorders.

## Introduction

The innate immune system exploits a large array of pattern-recognition receptors to detect pathogen- or danger- associated molecules to initiate a protective immune response (Janeway and Medzhitov, 2002). A subset of immune sensor proteins belong to the Nod-like receptor protein (NLR) family and function as sensors for the inflammasome complex, a conserved macro-molecular platform that governs inflammation driven by the interleukin-1 family of cytokines. The mammalian inflammasome complex minimally consists of an NLR sensor, the adaptor protein ASC and the effector inflammatory caspase, caspase-1 (Davis et al., 2011; Martinon et al., 2002; Tschopp et al., 2003). Upon ligand engagement by NLR sensors, the inflammasome complex initiates a distinct form of inflammatory cell death known as ‘pyroptosis‘. The distinguishing features and mediators of inflammasome-driven pyroptosis have been defined at the molecular level and include: ‘prionoid-like’ assembly of ASC (Lu et al., 2014), proteolytic activation of caspase-1, processing of pro-IL-1B and pro-IL-18 into their respective bioactive mature forms, extracellular secretion of mature IL-1B and IL-18 (Martinon et al., 2002), and lytic cell death following GSDMD-mediated membrane disruption (Kayagaki et al., 2015; Shi et al., 2015). In concert with other innate immune pathways, the inflammasome plays an important role in immune defense against bacterial and viral infections (Lamkanfi and Dixit, 2011; Lamkanfi et al., 2007), as demonstrated in the increased pathogen susceptibility in a variety of inflammasome knockout animal models.

In addition to robust, accurate and sensitive sensing of infection- or danger-related triggers, NLR proteins must avoid spontaneous and aberrant activation of ‘sterile inflammation‘, which can lead to host tissue damage. Recent work has revealed that the inflammasome employs a network of post-transcriptional and post-translational regulatory ‘checkpoints’ to guard against aberrant activation, including direct NLR modification and obligate regulatory factors (Guo et al., 2016; Kim et al., 2015; Qu et al., 2012; Shoham et al., 2003; Spalinger et al., 2016; Stutz et al., 2017; Xu et al., 2014). The *in vivo* importance of this regulatory network in preventing pathological inflammation is illustrated by a group of Mendelian genetic diseases caused by mutations in inflammasome components. Most of the inflammasome-related disorders are caused by germline mutations in NLR sensor proteins or their negative regulators, resulting in unprovoked periodic fever and macrophage/monocyte activation caused by constitutive and persistent inflammasome activation (Kastner et al., 2010; Moghaddas and Masters, 2015). In addition, dysregulation of NLR-driven inflammasome response has also been implicated in non-Mendelian diseases such as cancer, auto-immune and neuro-degenerative diseases (Davis et al., 2011; Venegas et al., 2017). Hence, there is an important need to gain a fuller molecular understanding of how various NLR proteins are maintained in the inactive state without compromising their ability to readily activate inflammation upon ligand engagement.

We and others have recently characterized a unique member of the NLR family, NLRP1. Although NLRP1 was one of the first NLR proteins shown to function as an inflammasome sensor, its cognate ligands and endogenous regulation remain poorly understood in human cells. NLRP1 differs from other known NLR sensor proteins in several aspects. Patients who have germline mutations in *NLRP1* all experience early-onset epithelial hyperkeratosis/dykeratosis, particularly on palmoplantar skin and in the eyes, while classical signs of fever or auto-inflammation that define other inflammasome activation diseases are variable (Grandemange et al., 2017; Zhong et al., 2016). This is partially explained by the high level of expression of NLRP1 in squamous epithelia as compared to other NLRs. In fact, NLRP1 is likely the only inflammasome sensor expressed in uninflamed primary human skin (Sand et al., 2018; Zhong et al., 2016). On the molecular level, human NLRP1 harbors an atypical pyrin domain (PYD) that is required for NLRP1 auto-inhibition, in contrast to PYDs of other NLRs such as NLRP3, AIM2 and MEFV (Finger et al., 2012; Zhong et al., 2016). NLRP1 assembles the inflammasome adaptor protein ASC via its CARD in a non-canonical pathway that requires auto-proteolysis within a domain of unknown function termed FIIND (D’Osualdo et al., 2011; Finger et al., 2012; Zhong et al., 2016). Although specific pathogen-derived triggers have been identified for certain rodent Nlrp1 alleles (Chavarria-Smith and Vance, 2013; Cirelli et al., 2014; Ewald et al., 2014), no specific agonists or dedicated regulatory co-factors have been reported for human NLRP1.

Here we report the identification of dipeptidyl peptidase, DPP9 as an evolutionarily conserved, endogenous interacting partner and inhibitor of NLRP1 in primary human and mouse cells. Inhibition of DPP9 via small molecule inhibitors of its peptidase activity, targeted mutations of its catalytic site and genetic deletion act as potent triggers for NLRP1-dependent inflammatory death, which proceeds via NLRP1 oligomerization, ASC speck assembly and IL-1 cleavage in a range of primary cell types. Mechanistically we identify NLRP1-FIIND as a self-sufficient DPP9 binding domain whose disruption by a patient-derived point mutation leads to spontaneous NLRP1 inflammasome activation without impacting NLRP1 auto-proteolysis. This likely explains the persistent sterile inflammation seen in in the auto-inflammatory/auto-immune syndrome, NAIAD. Our findings highlight an unprecedented peptidase-based regulatory checkpoint for an inflammasome sensor that could be of broad relevance in human immunity and inflammatory diseases.

## Results

### Identification of DPP9 as a novel binding partner of full-length, auto-inhibited NLRP1

To search for novel proteins that are involved in NLRP1 regulation, we took advantage of our previous observation that full-length NLRP1 is minimally active when expressed in 293T cells whereas the NLRP1 auto-proteolytic fragment (a.a. 1214-1474) is constitutively active (Finger et al., 2012; Zhong et al., 2016). We thus hypothesized that 293T cells might express additional unknown factors that interact with the regulatory domains of NLRP1 (PYD, NACHT, LRR and FIIND) to help maintain NLRP1 self-inhibition in the absence of its cognate ligands. To identify such factors, we performed immuno-precipitation (IP) of FLAG-tagged full-length NLRP1 expressed in 293T cells, with the constitutively active fragment as a negative control (Figure 1A). Direct protein staining of the IP eluates following SDS-PAGE revealed a prominent band at ~100 kDa that co-purified only with full length NLRP1, but not a.a. 1213-1474 (Figure 1B). Quantitative mass spec by isobaric labeling of the FLAG IP eluates identified this candidate interacting protein as the long isoform of dipeptidyl peptidase, DPP9 (Uniprot Accession Q86TI2-2) (Figure 1C). It was amongst the most enriched proteins that associated with full length NLRP1, but not with vector transfected cells, or cells expressing NLRP1 a.a. 1213-1474 or a.a. 1213-1373 (fold change >16, Figure S1A, B). Unlike other enriched proteins, DPP9 had not been observed as a common contaminant in IP-mass spec experiments (Figure S1B). Human DPP9 is a dipeptidyl dipeptidase of the DPP-IV family with broad functions in immune regulation, growth factor signaling, adipocyte differentiation and cellular metabolism (Gall et al., 2013; Justa-Schuch et al., 2016; Kim et al., 2017). It shares a similar domain structure with other family members consisting of an N-terminal β-barrel (DPP-IV N) and a C-terminal S9 hydrolase domain (Figure 1D). Out of all the DPP-IV family members, only DPP9 was detected as a specific interacting protein with NLRP1 (Figure 1E). Using a validated antibody, endogenous DPP9 was detected by western blot in the full length NLRP1 IP eluate, but not in four control IP eluates (Figure S1C,D). We further established that both DPP9 splice isoforms, DPP9L and DPP9S coimmunoprecipitate with full length NLRP1 (Figure 1F, lane 6 and 7), but not DPP9L lacking the hydrolase domain (Figure 1F, lane 6 vs. lane 8). In addition, a related inflammasome sensor NLRP3 did not interact with endogenous DPP9 (Figure 1G, lane 4, vs lane 5). These results suggest that DPP9 is a specific NLRP1 interacting partner and a candidate regulatory factor maintaining full-length NLRP1 self-inhibition.

**Figure 1.**
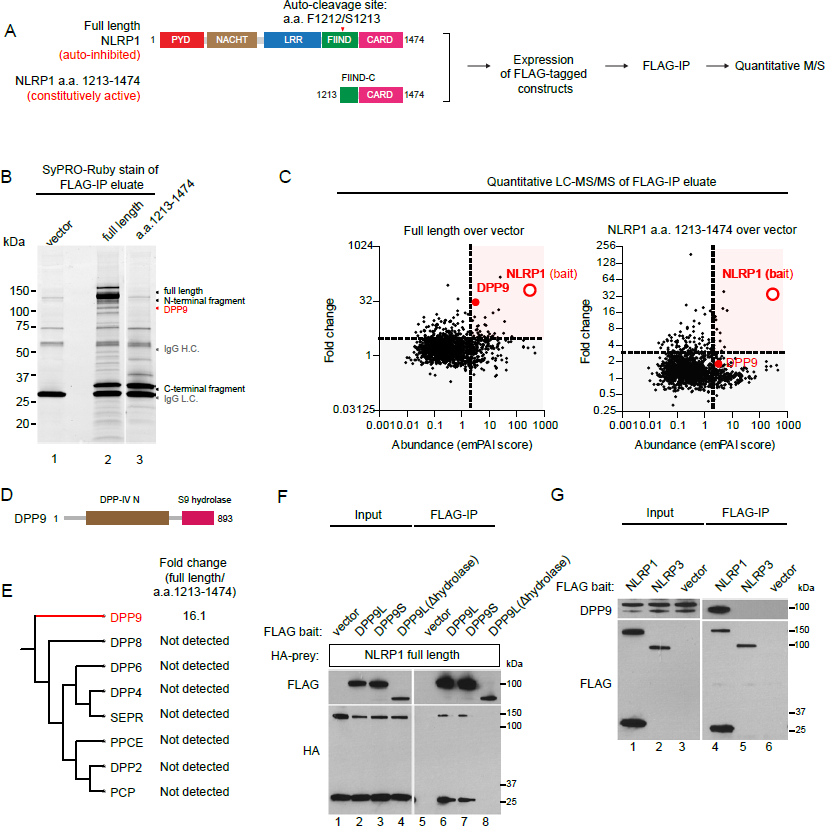
Identification of dipeptidyl peptidase DPP9 as a specific interacting protein for full length, auto-inhibited NLRP1. A. Proteomics-based strategy to identify NLRP1 interacting proteins. 293T cells were transfected with NLRP1 full length and NLRP1 a.a. 1213-1474 expressing constructs, expanded and harvested 4 days post transfection. Approximately 10^8^ cells were used per immunoprecipitation. B. Direct staining of NLRP1 interacting proteins after immuno-purification from 293T cells. C. Quantitative comparison of proteins that specifically associate with full length NLRP1 vs. constitutively active, auto-proteolytic NLRP1 fragment, a.a. 1213-1474. Fold change cut-off=3; Abundance (emPAI) cut-off=5 D. Domain structure of DPP9. E. Primary sequence alignment of related human dipeptidases and their fold enrichment in the NLRP1 FLAG-IP eluates vs. a.a. 1213-1474. F. The peptidase domain for DPP9 is required for NLRP1 binding. 293T cells were transfected with the indicated constructs and harvested 2 days post-transfection. 2 million cells were used for anti-FLAG immunoprecipitation. G. NLRP3 does not associate with DPP9. Immunoprecipitation was performed as in **Related to Figure S1**

### DPP9 inhibition triggers NLRP1 oligomerization, ASC speck formation and inflammasome activation

The chemical biology of DPP-IV family of peptidases has been extensively investigated due to the prominence of DPP4 as an effective anti-diabetic drug target (Pratley and Salsali, 2007), and a number of small molecule inhibitors for DPP9 have been developed (Yazbeck et al., 2009). In addition, DPP8/9 inhibitors have recently been suggested to activate an atypical form of pyroptotic cell death that does not involve ASC or IL-1 cleavage, though the mechanism remains unclear (Okondo et al., 2017; Taabazuing et al., 2017). This prompted to us to consider if the enzymatic inhibition of DPP9 could directly activate the NLRP1 inflammasome. To test this, we reconstituted the NLRP1 inflammasome in a 293T reporter cell line that stably expressed GFP-tagged inflammasome adaptor, ASC (293T-ASC-GFP). When NLRP1-expressing 293T-ASCGFP cells were treated with a pan-DPP-IV inhibitor, Talabostat (0.3 μM) or a specific DPP8/9 inhibitor 1G244 (10 μM), more than 70% of the cells formed large ASC-GFP specks that represented activated, assembled inflammasome complexes (Figure 2A, B). This effect was not observed in the absence of NLRP1 (Figure 2B, white bars) or in ASCGFP cells reconstituted with NLRP3 (Figure 2B, gray bars). We further confirmed the presence of ASC polymerization via DSS crosslinking followed by Western blot. Talabostat triggered a significant increase in ASC-GFP oligomers (Figure 2C, lane 5 vs 4), while a specific DPP4 inhibitor, sitagliptin, which has no cross-reactivity to DPP8/9 (Green et al., 2015), failed to nucleate ASC-GFP specks in NLRP1 inflammasome reconstituted cells (Figure 2B) or induce ASC-GFP polymerization (Figure 2C, lane 6 vs. lane 3). Notably, Talabostat or 1G244 did not enhance constitutive inflammasome activation by a known gain-of-function NLRP1 pyrin domain (PYD) mutation (p. M77T) found in patients with multiple self-healing palmoplantar carcinoma (MSPC) (Figure 2B, pink bars; Figure 2C, lanes 7-9). Taken together, these results demonstrate that enzymatic inhibition of DPP9 specifically activates the reconstituted NLRP1 inflammasome and induces polymerization of inflammasome adaptor protein ASC, likely independently of the PYD. These results also suggest that the reported, pro-pyroptotic effect of Talabostat might occur via NLRP1, rather than acting downstream of ASC polymerization as suggested (Okondo et al., 2017).

**Figure 2.**
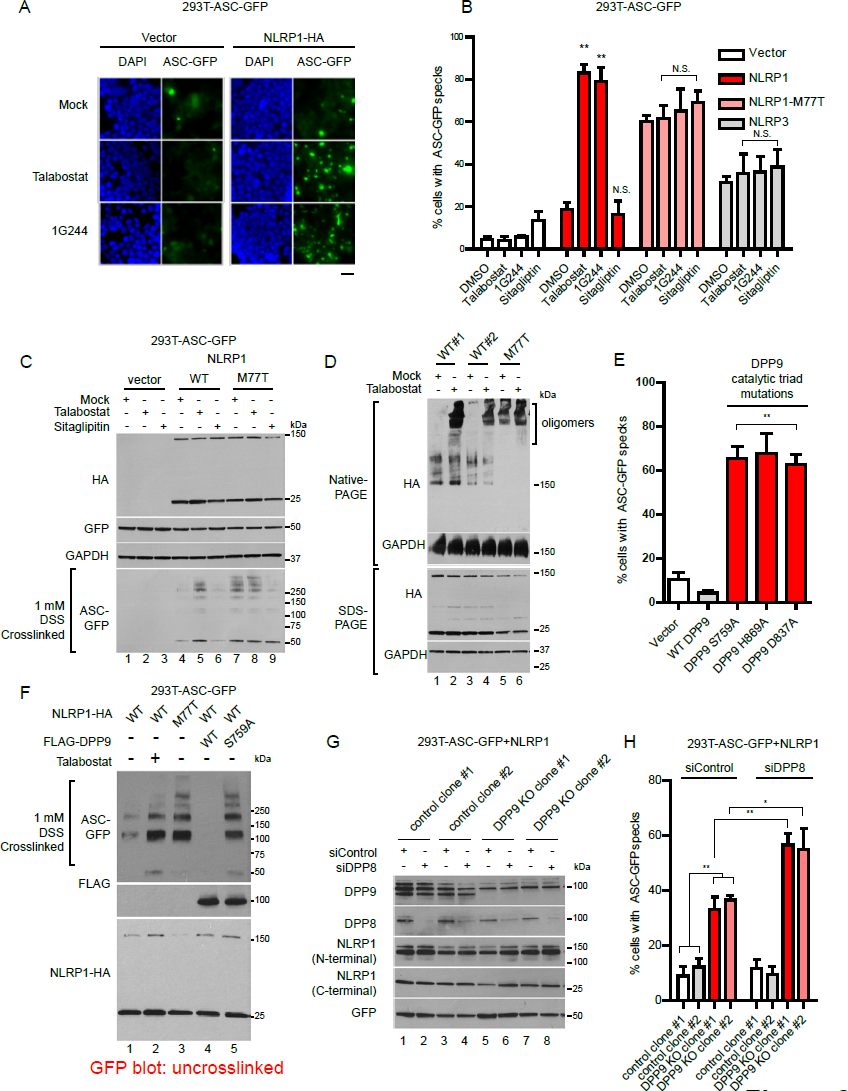
DPP9 inhibition or deletion triggers rapid NLRP1 oligomerization and ASC inflammasome assembly. A. DPP8/9 inhibitors cause ASC-GFP speck formation in the presence of NLRP1. 293T-ASC-GFP cells were transfected NLRP1 expressing plasmids at a ratio of 1 μg/5x10^5^ cells. Transfected cells were treated with Talabostat (0.2 μM) and 1G244 (10 μM) for 16 hours before GFP imaging. Scale bar=20 μm. B. DPP8/9 inhibition does not activate the NLRP3 inflammasome or enhance a NLRP1 pyrin-domain (PYD) mutant, p. M77T. 293T-ASC-GFP cells were transfected and treated as in A. C. Talabostat leads to ASC oligomerization independently of DPP4. 293T-ASC-GFP cells were transfected and treated as in A. Cell pellets were lysed in 1xTBS buffer with 1% NP-40. Insoluble pellets were subjected to crosslinking with 1 mM DSS for 15 mins at 37 °C and solubilized in 1xTBS with 1% SDS. D. DPP8/9 inhibition by Talabostat causes NLRP1 self-oligomerization. 293T cells were transfected with the respective constructs at a ratio of 2 μg/5x10^5^ cells and treated with Talabostat (2 μM) two days after transfection for 16 hours. E. Alanine mutations in the DPP9 catalytic triad dominantly activate the NLRP1-ASC inflammasome. 293T-ASC-GFP cells were co-transfected with NLRP1 and the respective DPP9 constructs and imaged two days after transfection. F. Opposing roles of wild-type DPP9 and S759A mutant in mediating ASC-GFP oligomerization in the presence of NLRP1. DSS crosslinking was performed as in C. G. Validation of CRISPR/Cas9-mediated deletion of DPP9 and subsequent knockdown of DPP8. Cells were harvested and lysed in 1xTBS buffer with 1% NP-40 4 days after siRNA transfection. H. DPP9 deletion activates the NLRP1 inflammasome with partial compensation by DPP8. 293T-ASC-GFP cells were treated with control or siRNAs against *DPP8* for 3 days before NLRP1 transfection. **Related to Figure S2**

To test this hypothesis and further probe the mechanism of how DPP9 inhibition might directly activate NLRP1 as an inflammasome sensor, we examined the monomer-to-oligomer transition of NLRP1. Previously we established that NLRP1 activation requires an obligatory oligomerization step that is regulated by its N-terminal domains including PYD, NACHT and LRR. Human germline mutations in these domains, such as p.M77T (Figure 2B,C) result in constitutive NLRP1 oligomer formation and cause aberrant inflammasome activation seen in patients (Zhong et al., 2016). When NLRP1 expressing cells were treated with Talabostat, a substantial amount of NLRP1 underwent oligomerization into ~1 MDa high molecular weight species as detected by Blue-Native PAGE, while the total level of NLRP1 remained unchanged (Figure 2D, SDS-PAGE, lane 2 vs.1 and lane 4 vs. 3). As this occurred in the absence of ASC, these results suggest that DPP9 exerts its inhibitory effect directly on NLRP1 by preventing its transition from inactive monomers to active oligomers. NLRP1 auto-cleavage generates an active C-terminal fragment, a.a. 1213-1474 (Figure 1A). We noted that this creates a potential DPP9 processing site with a proline residue at the P2 position. In the mass spec analysis of full-length NLRP1 immuno-purified from 293T cells, we readily detected tryptic peptides spanning and beginning at the auto-cleavage junction, which correspond to the uncleaved and auto-cleaved forms of NLRP1. However, we did not observe peptides that were consistent with DPP9 processing (i.e. starting at a.a. L1215), suggesting that NLRP1 might not be a direct *enzymatic* substrate of DPP9 (Figure S2A). We also confirmed that Talabostat did not engage NLRP1 itself using a cellular thermoshift assay (Figure S2B).

To rule out potential off-target effects of these chemical inhibitors, we exploited previous biochemical findings that DPP-IV family enzymes, including DPP9 are obligate dimers. As a result, enzymatic dead mutants can function as dominant negative inhibitors upon overexpression, likely by sequestering the wild-type subunits (Tang et al., 2011). The conserved ‘catalytic triad’ residues (S759, H869 and D837) were individually mutated to alanine to generate three dominant negative DPP9 constructs. When expressed in NLRP1 reconstituted 293T-ASC-GFP reporter cells, each of the catalytic triad mutant resulted in >60% of the cells forming ASC-GFP specks similar to Talabostat (Figure 2E, red bars), while wild-type DPP9 showed the opposite effect- by suppressing the low basal level of ASC-GFP speck formation in wild-type NLRP1 expressing cells (Figure 2E, gray bar). This was corroborated by direct visualization of ASC-GFP polymerization after covalent crosslinking with DSS (Figure 2F, lanes 4 and 5 vs lane 1). These results offer orthogonal evidence that inhibition of the endogenous DPP9 enzyme triggers NLRP1 activation. To prove genetically that DPP9 is required to maintain NLRP1 in the inactive state, we generated *DPP9* KO 293T-ASC-GFP cells using CRISPR/Cas9 (Figure 2G, lanes 1-4 vs. 5-8). In two independent clones, we observed a significantly higher percentage of cells forming ASC-GFP specks upon NLRP1 expression (Figure 2H, pink bars), albeit at a lower level than in Talabostat-treated cells or cells transfected with dominant negative DPP9 mutants (Figure 2B, 2E). We speculate that the clonal selection of KO cells over 3 weeks might allow for genetic compensation not seen in contexts of acute DPP9 inhibition. To test this, we knocked down the orthologous enzyme DPP8 (Figure 2G, lanes 1, 3, 5, 7 vs. lanes 2, 4, 6, 8), which has overlapping substrate preference with DPP9 (Wilson et al., 2013) and is also inhibited by Talabostat and 1G244. Partial knockdown of endogenous DPP8 caused a further increase in the percentage of cells forming ASC-GFP specks (Figure 2H, red bars). Taken together with data from acute DPP9 inhibition, these findings establish that DPP9 functions as a novel and cognate inhibitor of NLRP1. In the absence of any trigger, DPP9 enzymatic activity acts as a ‘brake’ against aberrant NLRP1-inflammasome activation.

### DPP9 inhibition leads to NLRP1-dependent pyroptosis via ASC oligomerization, caspase-1 activation and mature IL-1 secretion

We next investigated the effect of DPP9 on NLRP1 in two primary human cell types that are of direct relevance to NLRP1-associated auto-inflammatory diseases: skin keratinocytes and freshly isolated peripheral blood mononuclear cells (PBMCs). While PBMCs express a number NLR sensors including NLRP1, we and others have recently established that NLRP1, rather than NLRP3 is the most prominent, if not the only inflammasome sensor expressed in resting human primary and immortalized, non-transformed keratinocytes (Zhong et al., 2016). This provides a unique system to directly and specifically investigate the regulation of NLRP1 without impinging upon other NLR sensors such as NLRP3. Both keratinocytes and PBMCs secreted large amounts of mature IL-1B into the culture medium upon Talabostat treatment (3-30 μM) (Figure 3A, >1000 fold). In PBMCs isolated from two out of three donors, the amount of IL-1B secretion after Talabostat treatment exceeded that elicited by 1 μg/ml of LPS (Figure S3A, gray bars). Prior LPS stimulation further enhanced the effect of Talabostat (Figure S3A, red bars). In addition, direct transfection of immortalized keratinocytes with a catalytically inactive mutant of DPP9, S759A induced IL-1B secretion to the same extent as the constitutively active fragment of NLRP1, a.a. 1213-1474 (Figure 3F). A broader panel of chemical inhibitors confirmed that all DPP8/9 inhibitors were able to cause IL-1B secretion in keratinocytes, with a positive correlation between the IC50 against DPP8/9 and degree of inflammasome activation (Figure S3H). These results suggest that DPP9 is a potent inducer of IL-1B secretion in primary human cells. We further used Luminex to characterize in greater detail the chemokine/cytokine signature elicited by DPP9 inhibition. In both keratinocytes and PBMCs, IL-1B is the most significantly enriched cytokine following Talabostat exposure. In the case of keratinocytes, Talabostat led to a chemokine/cytokine profile that closely mimics that of primary keratinocytes derived from MSPC and FKLC patients with germline gain-of-function mutations in NLRP1 (Figure 3C and S3B) (Zhong et al., 2016), demonstrating that DPP9 inhibition has a strikingly similar effect as constitutive NLRP1 activation. In PBMCs, Talabostat alone led to IL-1B and IL-1A secretion (Figure 3D) without prior priming; it also caused the secretion of other inflammatory cytokines such as IL-6, TNF-α, GM-CSF, MIP-1a/b and IL-8 (Figure S3C), which are suggestive of monocyte activation (Figure S3C). When PBMCs were pre-stimulated with LPS, DPP9 inhibition by Talabostat led to further increase in IL-1B secretion (Figure S3D), while the only other cytokine that demonstrated a synergistic effect (>2 fold increase) was IL-1A, whose secretion also requires inflammasome activation in certain contexts (Gross et al., 2012). These data support the notion that DPP9 inhibition might be a highly specific trigger for an IL-1 driven inflammatory response in diverse primary cell types.

**Figure 3.**
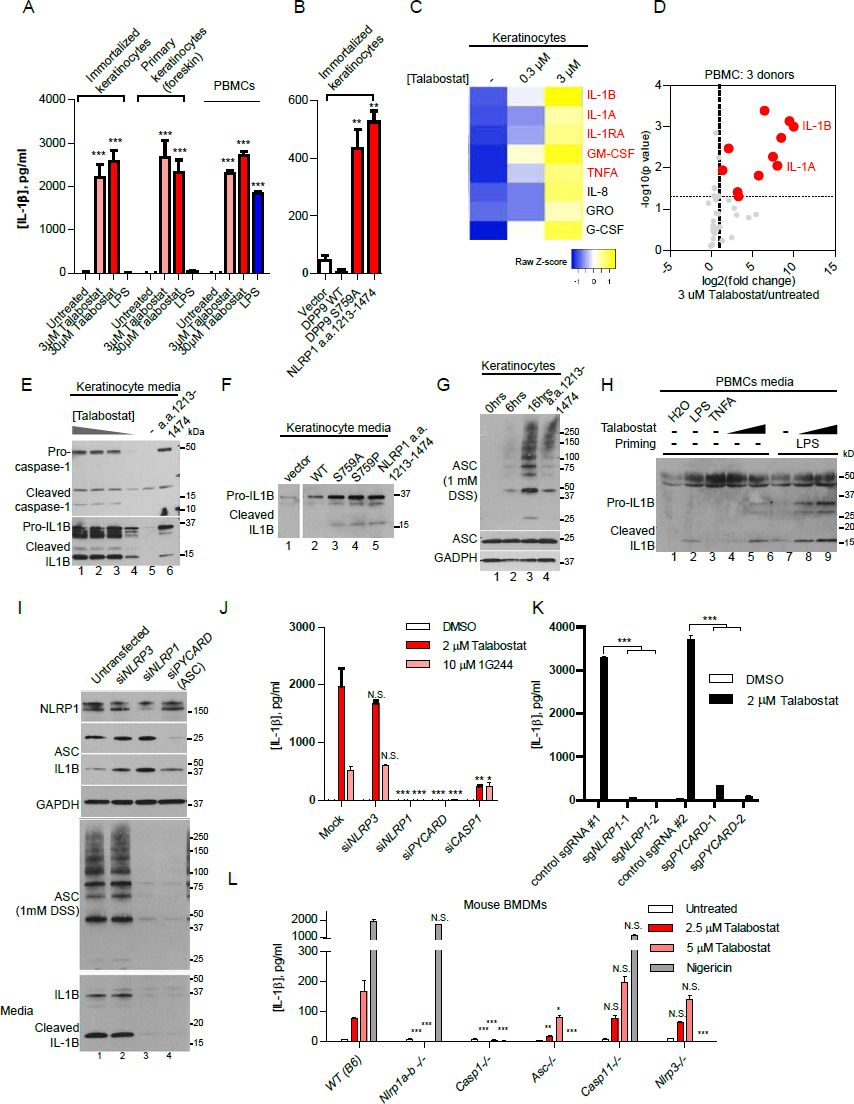
Primary human cells undergo NLRP1- and ASC-dependent pyroptotic cell death and mature IL-1B secretion upon DPP9 inhibition. A. IL-1B secretion from primary human keratinocytes and PBMCs upon Talabostat (2 μM) treatment. B. Human keratinocytes secrete IL-1B upon DPP9 S759A expression. Human keratinocytes were transfected with the respective plasmids with a ratio of 0.5 μg/ 5 x10^5^ cells. Conditioned media were harvested 24 hours post transfection. C. Cykokine/chemokine response of keratinocytes to Talabostat is highly similar to MSPC/FKLC patient-derived keratinocytes harboring gain-of-function NLRP1 mutations. Luminex array was performed on conditioned media of Talabostat-treated keratinocytes. Cytokines/chemokines that were also enriched in MSPC/FKLC patient-derived primary keratinocytes are shown in red. D. Cytokine/chemokine analysis of PBMCs treated with 2 μM Talabostat. Luminex array was performed on conditioned media of Talabostat-treated PBMCs isolated from three donors. P-values were calculated based on Student‘s t-tests after log transformation. E. DPP8/9 inhibition by Talabostat causes dose-dependent IL-1B processing. Cultured immortalized keratinocytes were treated with different 0.2 μM, 2 μM, 20 μM and 200 μM Talabostat for 24 hours. Conditioned media was concentrated 10 times for SDS-PAGE. F. DPP8/9 inhibition by S759A and S759P leads to IL-1B processing and secretion. Keratinocytes were transfected with the DPP9 expressing constructs. Conditioned media was harvested 24 hours post-transfection. G. DPP8/9 inhibition by Talabostat causes endogenous ASC oligomerization. DSS crosslinking was performed as in Figure 2C. H. DPP8/9 inhibition by Talabostat leads to IL-1B processing and secretion in PBMCs. Conditioned media from PBMC (Donor 3) was used for SDS-PAGE without prior concentration. I. NLRP1 and ASC knockdown abrogates Talabostat-induced ASC oligomerization and IL-1B processing. Immortalized keratinocytes were treated with 2 μM Talabostat for 24 hours three days after siRNA incubation. Conditioned media was concentrated 10 times before SDS-PAGE. J. Genetic requirement of *NLRP1*, *ASC* and *CASP1*, but not NLRP3 in the effect of DPP8/9 inhibition. Immortalized keratinocytes treated with siRNAs and DPP8/9 inhibitors as in J. Conditioned media was diluted 1:5 before IL-1B ELISA. K. CRISPR/Cas9-mediated deletion of NLRP1 and ASC blocks Talabostat-induced pyroptosis. L. Dissection of the genetic requirement for inflammasome activation upon DPP8/9 inhibition in mouse bone marrow derived macrophages (BMDMs). Murine BMDMs from the indicated genotypes were primed with LPS (200 ng/ml), then treated with the indicated concentrations of Talabostat for 24 hours. **Related to Figure S3**

Talabostat and 1G244 were recently shown to cause caspase-1 dependent pyroptosis in human cancer cells without ASC and IL-1B cleavage. These characteristics are somewhat inconsistent with the involvement of NLRP1, which requires ASC to activate the inflammasome response in human cells (Zhong et al., 2016). It is noteworthy that some commonly used cancer cell lines have markedly different regulatory mechanisms of inflammasome activation from primary cells (Gaidt et al., 2017; Gaidt et al., 2016). To clarify the mechanism of DPP9 inhibition in disease-relevant primary human cells, we examined the molecular hallmarks of ‘canonical’ inflammasome activation, i.e. lytic cell death, IL-1B cleavage and ASC polymerization. Talabostat induced rapid swelling of primary and immortalized keratinocytes (Figure S3E) and significant cell death in PBMCs (Figure S3G). This was accompanied by dose-dependent cleavage of pro-caspase-1 and pro-IL-1B into their respective mature, secreted forms by both keratinocytes and freshly isolated PBMCs (Figure 3E and H). Furthermore, Talabostat caused time-dependent ASC polymerization and speck formation in keratinocytes (Figure 3G, S3F). We similarly observed robust IL-1B cleavage and secretion in primary human keratinocytes transfected with two distinct dominant negative DPP9 mutants (Figure 3F). Hence, in a variety of primary human cells, DPP9 inhibition induces a canonical inflammasome response involving lytic cell death, caspase-1 activation, IL-1B processing and ASC polymerization.

Given our biochemical findings on reconstituted NLRP1 inflammasome, we further postulated that DPP9 inhibition specifically acts upstream of and must require NLRP1 and ASC for pyroptosis induction. To test this genetically, immortalized keratinocytes were pre-treated with siRNAs for three days before DPP9 inhibition for 16 hours. siRNAs against *NLRP3*, which is not expressed in keratinocytes (RNAseq FKPM<1, EnCODE) was included as a negative control. Effective protein depletion (>70%) was observed at Day 4 (Figure 3I, top). In contrast to untransfected and si*NLRP3* treated keratinocytes, NLRP1-depleted keratinocytes failed to undergo ASC polymerization as measured by DSS crosslinking (Figure 3I, middle, lane 3 and 4 vs. 1 and 2). In addition, NLRP1, ASC and caspase-1 depletion abrogated IL-1B processing and secretion (Figure 3I, bottom, lane 3 and 4 vs. 1 and 2, Figure S3F), in comparison to untransfected and si*NLRP3-*treated controls. Similar results were obtained using 1G244, a more specific but less potent DPP8/9 inhibitor (Figure 3J). Furthermore, CRISPR/Cas9-mediated disruption of the *NLRP1* and *PYCARD* (encoding ASC) loci abrogated IL-1B secretion following Talabostat treatment (Figure 3J). Together, these results provide genetic evidence that DPP9 inhibition specifically activates the NLRP1 inflammasome and IL-1 driven inflammation in primary, untransformed human cells in a manner that requires ASC and caspase-1. We further took advantage of a suite of inflammasome KO murine models to investigate if DPP8/9 inhibition plays an evolutionarily conserved role as an NLRP1 inflammasome inhibitor, despite the divergence of NLRP1 genetic and protein structure. Using primary bone marrow derived macrophages (BMDMs), we found that genetic ablation of all *Nlrp1* isoforms (including *Nlrp1a,b and c*) and *Casp1* blocked IL-1B secretion following Talabostat exposure, while Nlrp3 and Casp11 KO BMDMs did not differ significantly from wild-type (B6) controls. Notably, deletion of *Pycard/Asc* also significantly reduced IL-1B secretion with at a lower dose of Talabostat (>5 fold, 2.5 μM Talabostat), in agreement with our findings using human cells (Figure 3L, red bars). The effect of *Pycard/Asc* KO was significantly less pronounced at a higher dose of Talabostat (~2 fold, 10 μM Talabostat, pink bars), suggesting that murine Nlrp1 might be able to trigger inflammasome activation without ASC as previously suggested (Guey et al., 2014; Van Opdenbosch et al., 2014), but only in response to strong agonist triggers. As an additional control, *Nlrp1(a-c)* KO cells responded similarly to nigericin, a specific NLRP3 agonist, as wild-type cells (Figure 3L, gray bars). These findings provide further support that DPP8/9 inhibition specifically triggers NLRP1, but not other inflammasome complexes in an ASC- and caspase-1 dependent manner in both human and murine systems.

### A single FIIND mutation responsible for auto-inflammatory disorder NAIAD, disrupts DPP9 binding and leads to constitutive NLRP1 inflammasome activation

Our group and others have recently discovered that distinct germline, gain-of-function mutations in NLRP1 cause three allelic human Mendelian diseases, Multiple self-healing palmoplantar carcinoma (MSPC), familial keratosis lichenoid chronica (FLKC) and NLRP1-associated auto-inflammation with arthritis and dyskeratosis (NAIAD) (Figure 4A) (Grandemange et al., 2017; Zhong et al., 2016). These diseases differ considerably from other inflammasome disorders and, in the case of MSPC and FKLC, affect predominantly the squamous epithelia of the skin and the eyes. Mechanistically, we showed that most of the patient-derived mutations inactivate one of the NLRP1 auto-inhibitory domains, PYD and LRR (Figure 4A); however, one mutation p.P1214R found in an NAIAD patient is situated in the C-terminal auto-proteolytic fragment, a.a. 1213-1474, which is responsible for inflammasome activation (Figure 4A). Its pathogenic mechanism therefore remains unclear. To test if this mutation impacts the inhibitory effect DPP9 on NLRP1, we first carried out deletional mapping to identify the minimal DPP9 binding domain on NLRP1 (Figure 4B). NLRP1 shows a highly modular structure, and progressive removal of each NLRP1 domain revealed that PYD, NACHT, LRR and CARD were dispensable for DPP9 binding (Figure 4B, Figure 4C, lanes 6, 8, 9), while the intact FIIND domain (a.a. 986-1373) was required (Figure 4B, Figure 4C, lane 7 and 10). In agreement with previous data, we found that FIIND domain underwent auto-cleavage independently of other domains, but neither fragment alone was sufficient to bind DPP9 (Figure 4C, lane 10, Figure S1A, D). An intact FIIND in the absence of other domains was sufficient to interact with endogenous DPP9 by immunoprecipitation to a comparable extent as did wild-type NLRP1 (Figure 4D, lane 6 vs. lane 5 and lane 4). NLRP1-FIIND, whose function had remained elusive since its first description (D’Osualdo et al., 2011; Finger et al., 2012), can therefore be viewed as a necessary and self-sufficient DPP9 interacting domain.

**Figure 4.**
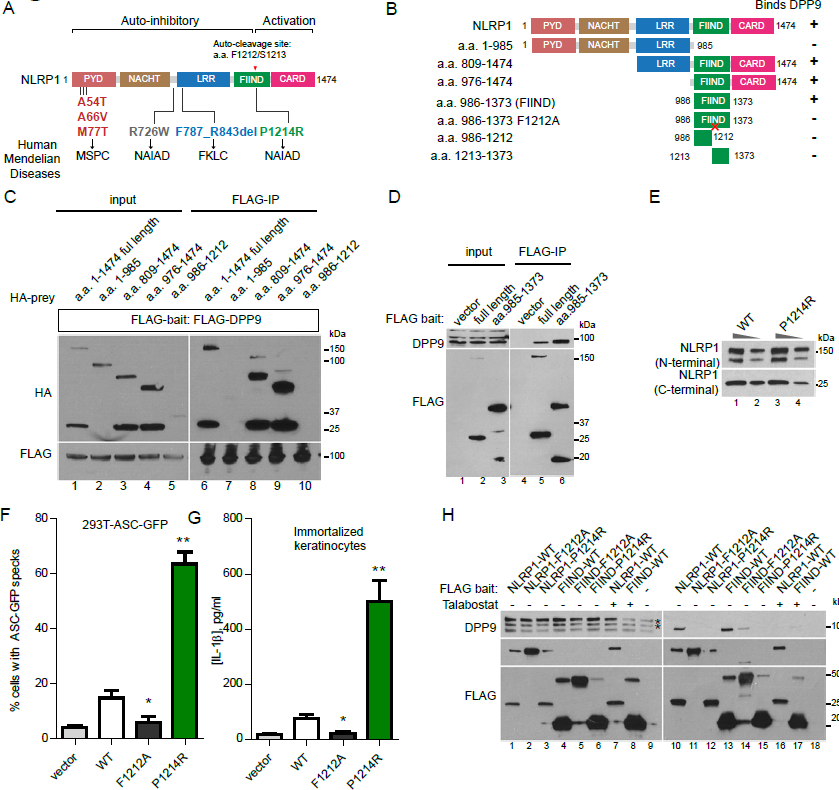
NLRP1 FIIND is a DPP9-interacting module and the disruption of NLRP1 FIIND-DPP9 binding explains the aberrant inflammasome activation in a human auto-inflammatory disorder. A. Overview of NLRP1 domain structure and all known pathogenic NLRP1 mutations in inherited human auto-inflammatory disorders. Auto-cleavage site within FIIND is shown in red. B. Summary of experimental results to identify the DPP9-binding domain in NLRP1. Binding was tested by anti-FLAG immunoprecipitation of the indicated NLRP1 fragments expressed in 293T cells followed by Western blot detection of endogenous DPP9. C. NLRP1 FIIND is required to bind DPP9. 293T cells were transfected with the indicated constructs and harvested 2 days post transfection. 2 million cells were used per anti-FLAG immunoprecipitation. D. NLRP1 FIIND is sufficient to bind DPP9. E. The NAIAD mutation P1214R does not affect FIIND auto-cleavage. Wild-type NLRP1 and NLRP1 p. P1214R were expressed in 293T cells. 10 μg or 2.5 μg of total cell lysate was used for SDS-PAGE. F. The NAIAD mutation P1214R causes ASC-GFP speck formation in a reporter cell line. 293T-ASC-GFP cells were transfected with wild-type NLRP1 or P1214R and imaged 24 hours post transfection. G. P1214R causes IL-1B secretion from human keratinocytes. Immortalized keratinocytes were transfected with wild-type NLRP1 or P1214R and imaged 24 hours post transfection. H. P1214R abrogates NLRP1-DPP9 binding, similar to Talabostat. Anti-FLAG immunoprecipitation was performed on 293T cells transfected with the indicated constructs as in C and D.

Although the NAIAD mutation, P1214R is found adjacent to the FIIND cleavage site, we did not observe any appreciable difference in its degree of auto-cleavage relative to wild-type NLRP1 when expressed in 293T cells (Figure 4E). However, it did potently abrogate NLRP1 auto-inhibition and led to constitutive inflammasome activation in both 293T-ASCGFP reporter cells (Figure 4F) and in immortalized human keratinocytes, as measured by IL-1B secretion (Figure 4G), in agreement with its causal role in NAIAD. Next, we tested the ability of P1214R to interact with endogenous DPP9 by co-IP. When expressed at similar levels, P1214R completely abrogated the ability of full length NLRP1 or NLRP1-FIIND to bind DPP9 (Figure 4G, lane 12 vs. 10, lane 15 vs 13), without affecting auto-cleavage. In addition, Talabostat treatment of wild-type NLRP1 expressing cells also abrogated NLRP1-DPP9 binding (Figure 4G, lane 16 vs. lane 10; lane 17 vs. lane 13). Hence, the loss of DPP9-NLRP1 interaction by a single point mutation in NLRP1-FIIND is sufficient to disrupt NLRP1 self-inhibition and lead to pathological auto-inflammation *in vivo* as observed in NAIAD patients.

## Discussion

Although NLRP1 was one of the first inflammasome sensors to be identified, its function in innate immunity and human inflammatory conditions has not been characterized in detail, presumably due to the lack of knowledge on its endogenous regulation and ligand specificity. Recently we and others have identified a group of Mendelian human inflammatory conditions caused by NLRP1 mutations, which demonstrate remarkable differences from other inflammasome disorders in terms of clinical presentation. Notably, all NLRP1 mutant patients display hyperkeratosis of the skin and other squamous epithelial organs. On the mechanistic level, all pathogenic NLRP1 mutations are gain-of-function and result in aberrant inflammasome activation in an ASC- and caspase-1-dependent manner, suggesting that the endogenous regulation of NLRP1 is critical in guarding against pathological auto-inflammation, particularly in non-hematologic organs such as the skin. In this report, we identify a protease, DPP9 as a specific interacting partner and inhibitor of NLRP1. DPP9 inhibition by small molecule compounds or dominant negative mutants potently and specifically activates the NLRP1 inflammasome in human keratinocytes and PBMCs. Our use of primary human cell types that are relevant to the NLRP1 mutant patient phenotypes was instrumental in deciphering its downstream signaling pathway. Surprisingly, in contrast to recently published data (Okondo et al., 2017; Taabazuing et al., 2017), we find that the inflammatory cell death elicited by DPP9 inhibition is dependent on NLRP1, ASC and caspase-1 in both human and murine cells and leads to robust cleavage and secretion of mature IL-1B.

Most importantly, we have delineated the mechanism by which DPP9 regulates NLRP1 activation. Our biochemical analysis shows that the DPP9 directly binds to NLRP1 via the FIIND domain, and thus maintains it in an inactive state. This assigns a new regulatory role to the FIIND domain, an auto-proteolytic domain with hitherto unknown function. This domain is only found in a handful of human proteins include CARD8 in which it is also followed by a C-terminal CARD domain. Like NLRP1, CARD8 is reported to participate in the inflammasome pathway and contribute to the pathogenesis of auto-inflammatory diseases (Cheung et al., 2017; D’Osualdo et al., 2011) so it is interesting to speculate if it is similarly bound and inhibited by an endogenous factor. Our work indicates that the FIIND of NLRP1 can be viewed as a self-sufficient DPP9 binding module. FIIND-DPP9 binding is critical for maintaining NLRP1 in the inactive monomeric state. Its disruption by the p. P1214R germline point mutation can explain the auto-inflammatory symptoms driven by constitutive activation of NLRP1 in NAIAD.

Taken together, we propose that NLRP1 acts as a sensor for a ‘homeostasis-altering molecular process’ (HAMP) (Liston and Masters, 2017) involving alterations of DPP9 enzymatic activity. DPP9 might be suppressed during infection by a foreign pathogen that is yet to be identified. This could be an integral part of the host immune response or elicited by a pathogen effector that seeks to manipulate host cell physiology to facilitate infection. In this case, alterations of DPP9 enzymatic activity would alert the host to a loss of cellular homeostasis, and this process therefore requires constant monitoring by a dedicated sensor- NLRP1. In the absence of any trigger, enzymatically active DPP9 ‘locks’ NLRP1 in the monomeric state to prevent aberrant activation. This is conceptually akin to the recently discovered mechanism by which another inflammasome sensor, Pyrin monitors Rho-GTPase for the presence of unusual post-translational modifications deposited by secreted bacterial toxins (Xu et al., 2014). We postulate that this HAMP detection system might allow human cells to mount a productive, inflammasome-driven immune response for rapid pathogen clearance. When its regulation is disrupted by germline mutations in NLRP1 or chemical/dominant-negative DPP9 inhibition, NLRP1 rapidly oligomerizes and assembles the inflammasome complex consisting of ASC and caspase-1 that initiates IL-1 driven inflammation. Our work provides the foundation for additional investigation of the biochemical nature of this novel immune regulatory mechanism and implicate DPP9 as a potential regulator in human inflammatory disorders.

## Materials and methods

### Cell culture

293Ts (laboratory stock) was cultured in DMEM supplemented with 10% fetal bovine serum (FBS) without antibiotics. Immortalized keratinocytes (N/TERT-1) were cultured in Keratinocyte Serum-free media with bovine pituary extract, EGF in the presence of 0.3mM CaCl2. PBMCs were obtained from donor using Ficoll and cultured in RPMI media with 10% FBS. Primary human keratinocytes were cultured using methods described by Rheinwald and Green (Rheinwald and Green, 1975). BMDM were prepared from the bone marrow of Nlrp1(abc)^−/−^ (Masters et al., 2012)), Casp1^−/−^ (Schott et al., 2004), Asc^−/−^ (Mariathasan et al., 2004), Casp11^−/−^ (Kayagaki et al., 2011), and Nlrp3^−/−^ (Martinon et al., 2006) mice, cultured in DMEM supplemented with 10% fetal bovine serum (FBS) and 10% L929-cell conditioned media for 6 days.

### Immunoprecipitation and mass spectrometry

Small scale immunoprecipitations were carried out by incubating overnight 0.5 mg total cell lysate prepared in 1xTBS-NP40 buffer (20mM Tris-HCl, 150 mM NaCl, 0.5% NP-40) and 10 μl of anti-FLAG-M2 agarose resin (Sigma-Aldrich) in 300~500 μl total volume. Bound proteins were eluted in 1x Laemmli‘s buffer at 95°C for 5 minutes.

For mass spectrometry, approximately 10^8^ transfected 293T cells were lysed in 3 ml 1xTBS-NP40 buffer and 100 μl anti-FLAG M2 agaorase beads were used for immunoprecipitation and directed subjected to trypsin digestion after washes in lysis buffer. Equal amount of peptides was taken for TMT isobaric tag (Thermo) labeling. Following labelling samples were combined, desalted and vacuum dried and subsequently re-suspended in 10mM Ammonia and using step gradient fractionated on C18 high pH reverse phase material using self-packed column (C-18 ReproSil-Pur Basic, Dr. Maisch, 10um) with 12, 25 and 50% of ACN in 10mM Ammonia Formate. Fractions were washed with 70% ACN with 0.1% formic acid and vacuum dried and subsequently analyzed using Easy nLC1000 (Thermo) chromatography system coupled with Orbitrap Fusion (Thermo). Each sample was separated in 120min gradient (0.1% Formic Acid in water and 99.9% Acetonitrile with 0.1% Formic Acid) using 50cm x 75m ID Easy-Spray column (C-18, 2m particles, Thermo). Data dependent mode was used with 3 sec cycle and Orbitrap analyser (ion targets and resolution OT-MS 4xE5 ions, resolution 60K, OTMS/MS 6E4 ions, resolution 15k). Peak lists were generated with Proteome Discoverer 2.2 software (Thermo) followed by searches with Mascot 2.6.1 (Matrix Science) against concatenated forward/decoy Human Uniprot database with following parameters: precursor mass tolerance (MS) 20ppm, OT-MS/MS 0.05 Da, 3 miss cleavages; Static modifications: Carboamidomethyl (C), TMT6plex. Variable modifications: Oxidation (M), Deamidated (NQ), Acetyl N-terminal protein. Forward/decoy searches were used for false discovery rate estimation (FDR 1%).

### Plasmid transfection and lentiviral transduction

All expression plasmids for transient expression was constructed based on the pCS2+ backbone and cloned using InFusion HD (Clonetech). All 293T transfection experiments were performed with Lipofectamine 2000 (ThermoFisher). Keratinocytes were transfected with Fugene HD (Promega). Lentiviral constructs were based on pCDH-puro (System Biosciences) and packaged with the third generation packaging plasmids.

### Chemical compounds

The small molecule inhibitors used are vildagliptin (MedChemExpress), saxagliptin (MedChemExpress), TC-E 5007 (Tocris), butabindide oxalate (MedChemExpress), Talabostat and (MedChemExpress), 1G244 (Santa Cruz Biotechnology), LPS (Ultrapure, Escherichia coli O111:B4, SigmaAldrich) and nigericin 10 uM (Invivogen, #tlrl-nig).

### Blue-Native and SDS-PAGE

Blue-Native PAGE was carried out using the Native-PAGE system (ThermoFisher) with 10-20 μg total cell lysate followed by dry transfer (TurboBlot, Bio-rad) and Western blot. SDS-PAGE was carried out using pre-cast TGX 4-20% gels (Bio-rad).

### CRISPR/Cas9 gene editing

DPP9 deletion in 293T-ASC-GFP cells was carried out using a co-editing and positive selection protocol adapted from by the Doyon lab (Agudelo et al., 2017) . Parental cells were co-transfected with plasmids (Plasmid #62988) encoding guide RNAs against *DPP9* and *ATP1A* (G2) in a ratio of 3:1. Clonal selection was carried out in the presence of 1 μM ouabain (SigmaAldrich). Lentiviral Cas9 and guide RNA plasmids (Addgene Plasmid #52962 and #52963) were used to create stable deletions of *NLRP1* and *PYCARD* in keratinocytes. The guide RNA target sequences are

*NLRP1*: CTATCAGCTGCTCTCGATAC, AGCCCGAGTGACATCGGTGG

*ASC*: CGCTAACGTGCTGCGCGACA, GCTAACGTGCTGCGCGACAT

*DPP9:* ATCCATGGCTGGTCCTACGG, TGTGTCGTAGGCCATCCAGA

**IL-1B ELISA and Luminex cytokine/chemokine array**

Human IL-1B was measured with Human IL-1B BD OptEIA ELISA kit. Mouse IL-1B ELISA was measured with R&D DY401 ELISA kit. Luminex cytokine/chemokine array was carried out using standard manufacturer-supplied protocol without modification.

### Measurement of cell death in PBMCs

After harvesting supernatants from PBMCs, the remaining cell pellets were used for quantification of cell death. Cells were washed in PBS, resuspended and incubated for 10 minutes in PBS containing 1:1000 LIVE/DEAD Fixable Green Dead Cell Stain (Thermo Fisher). The cells were then washed in staining buffer containing PBS, 0.2% (v/v) FBS and 2 mM EDTA. To identify immune cell lineages, a surface stain was performed for 20 minutes in Brilliant Stain Buffer (BD Biosciences) containing the following antibodies: CD4-BV786, CD8-Alexa Fluor 700, CD19-BUV496 (BD Biosciences), CD11c-BV421, CD123-BV650, CD56-BV711 (Biolegend) and CD14-Viogreen (Miltenyi Biotec). Samples were acquired using a FACSymphony flow cytometer (Becton-Dickinson) and analysed with FlowJo V10 (Flowjo LLC).

## Acknowledgements

We are grateful to all members of the B.R. laboratory for their support. B.R. is a fellow of the Branco Weiss Foundation and a recipient of the A*STAR Investigatorship and EMBO Young Investigator. B.R. and R.M.S. is supported by Core funding from IMCB and IMB Strategic Positioning Fund (SPF,BMRC, A*STAR), Young Investigator Grant YIG 2015 (BMRC, A*STAR), and NMRC MS-CETSA platform grant MOHIAFCAT2/004/2015. F.L.Z is supported by NMRC Young Investigator Grant NMRC/OFYIRG/0046/2017. S.L.M acknowledges funding from NHMRC grants (1142354 and 1099262), The Sylvia and Charles Viertel Foundation, HHMI-Wellcome International Research Scholarship and Glaxosmithkline.

## Author Contributions

F.L. Z. and B.R. conceptualized and designed the study. F.L.Z. performed all cell biology based experiments and analyzed the data with the help of K.R. and W.X̤ C.L., V.B.A and R.H. performed all Luminex and PBMC experiments and analyzed the data. R.M.S. carried out the mass spec experiment and analyzed the data. C.R.H. and C.H.Y performed BMDM experiments with the supervision of S.L.M. F.L.Z. wrote the manuscript with significant edits from B.R. and S.L.M.

---

**Figure S1.**
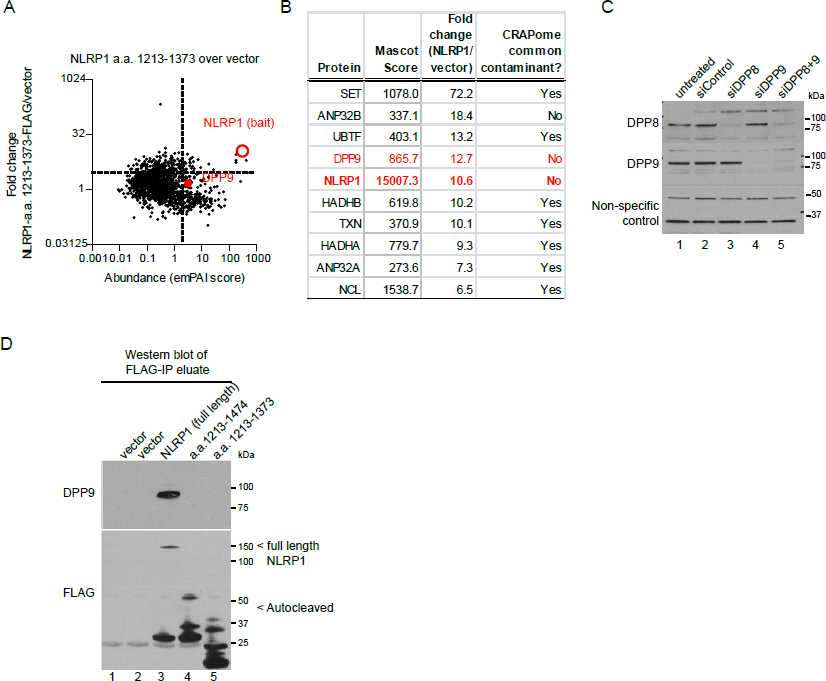
Additional evidence for the NLRP1-DPP9 interaction. A. DPP9 does not associate with NLRP1 a.a. 1213-1373, corresponding to the carboxy-terminal fragment of the FIIND domain. B. Top ten enriched proteins in the full length NLRP1 IP eluate. C. Validation of DPP8 and DPP9 antibodies in siRNA treated 293T cells. D. Direct Western detection of DPP9 in NLRP1 IP eluates vs. controls.

**Figure S2.**
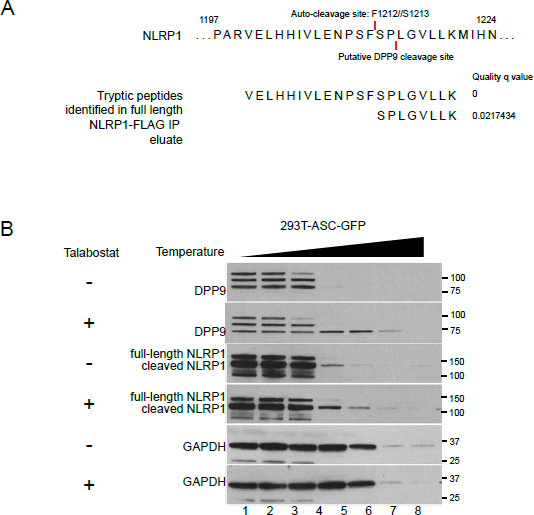
No evidence for direct NLRP1 processing by DPP9 or Talabostat binding to NLRP1. A. Lack of evidence for direct processing NLRP1 C-terminal fragments by DPP9 from MS tryptic peptide analysis. B. Lack of evidence for directly binding to NLRP1 by Talabostat. 293T-ASC-GFPNLRP1 cells were treated with 2 μM Talabostat for 16 hours and lysed by 3 rounds of freezing and thawing. 20 μl (2 μg/ul) clarified lysates were heated at a temperature gradient for 10 minutes, centrifuged at 16,000g for 10mins at room temperature. The supernatant was used for SDS-PAGE.

**Figure S3.**
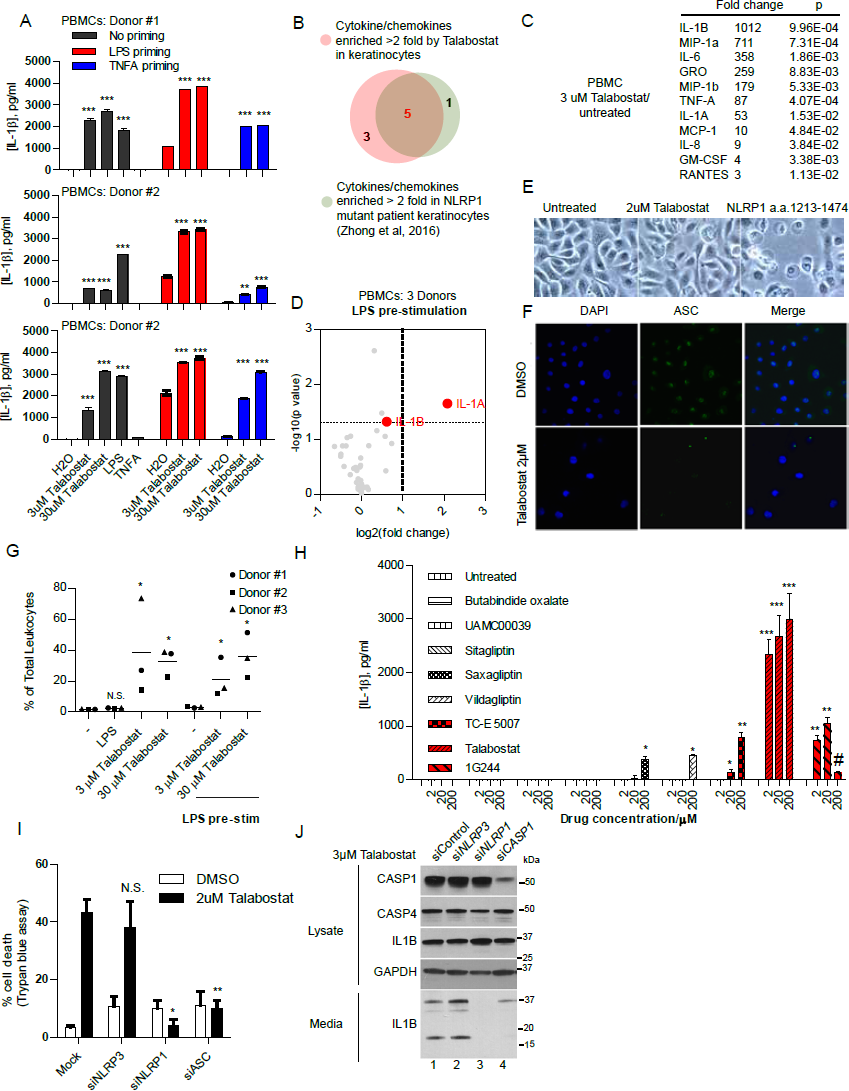
Additional evidence for the requirement of NLRP1, ASC and caspase-1 in the effect of DPP9 inhibition. A. Talabostat induces IL-1B secretion either alone or in cooperation with LPS-priming in primary human PBMCs. B. Overlap between Talabostat dependent cytokine/chemokine signature and MSPC/FKLC patient-derived keratinocytes. C. Luminex analysis of Talabostat-treated LPS-prestimulated PBMCs. D. List of cytokines/chemokines that are induced by Talabostat in PBMCs. E. Lytic cell death of Talabostat-treated keratinocytes. F. Talabostat induces ASC speck formation in immortalized human keratinocytes. G. Talabostat causes significant leukocyte cell death. H. The effect on IL-1B secretion by a panel of peptidase/protease inhibitors on immortalized keratinocytes. Compounds with known IC50 for DPP9<100 nM are colored red. I. Lytic death in keratinocytes upon Talabostat exposure requires NLRP1 and ASC, but not NLRP3. J. Caspase-1 knockdown abrogates the effect of Talabostat on IL-1B processing.

